# A marine probiotic treatment against the bacterial pathogen *Vibrio coralliilyticus* to improve the performance of Pacific (*Crassostrea gigas*) and Kumamoto (*C. sikamea*) oyster larvae

**DOI:** 10.1101/2022.05.09.491202

**Authors:** David Madison, Carla Schubiger, Spencer Lunda, Ryan S. Mueller, Chris Langdon

## Abstract

Oyster larvae reared in hatcheries on the U.S. West coast often experience severe *Vibrio coralliilyticus*-related mortalities early in their development. Current treatment options for these molluscs are either not available or feasible; however, for decades, probiotics have been successfully used in finfish and crustacean shellfish culture. Consequently, the objectives of this work were to 1) isolate marine bacteria from oysters and evaluate their protective activity against *Vibrio coralliilyticus* infection of Pacific oyster (*Crassostrea gigas*) larvae, and 2) to determine the long-term effects of probiotic additions on growth and metamorphosis of larval Pacific and Kumamoto oysters (*C. sikamea*). A combination of three probiotic strains applied once 24 hours post-fertilization was more effective in improving survival of larval *C. gigas* exposed to lethal concentrations of *V. coralliilyticus* strain RE22, compared with separate additions of individual probiotics. In addition, a single application of the probiotic combination to one-day-old larvae increased the larval metamorphosis success of *C. sikamea* and both the Midori and Myiagi stocks of *C. gigas*. These results suggest that probiotics are effective at preventing bacterial infections and can significantly improve performance of oyster larvae, using a single application early in their development.

**Highlights:** • A combination of marine bacteria improved survival of Pacific oyster larvae exposed to virulent *V. coralliilyticus*.

• Metamorphosis was increased after adding a single dose of probiotics to one-day-old larvae.

• Repetitive dosing after each water change was not superior to a single dose one day post egg-fertilization.

• A single dose of the probiotic combination resulted in larger size on day 12.

## 1. Introduction

In the USA, oyster farms are largely dependent on high-quality seed (“eyed” larvae) from oyster hatcheries. Oyster hatcheries, however, have periodically experienced severe larval losses during the past two decades, leading to seed shortages and supply disruptions (Elston et al., 2008; Richards et al., 2015). Initially, these losses were mainly attributed to ocean acidification (Barton et al., 2012; Gray et al., 2022), and currently many hatcheries employ sophisticated systems that measure and correct acidified incoming water. Unfortunately, these treatments have not entirely resolved the problem, suggesting other factors, such as pathogens, likely play a significant role in these losses and contribute to various sub-lethal effects on oyster health (Marques et al., 2006).

Bacteria from the genus *Vibrio* are omnipresent in marine and brackish waters as commensals, mutualists, or pathogens (Takemura et al., 2014). They are highly adaptable to changing ocean conditions, including increasing temperature, lower pH, and salinity, and can make up more than 50% of all detectable microbes during favorable conditions (Gilbert et al., 2012; Oh et al., 2009; Vezzulli et al., 2010). *Vibrio coralliilyticus* has been linked to massive die-offs of Pacific oyster (*Crassostrea gigas*) larvae in U.S. West coast hatcheries (Elston et al., 2008; Estes et al., 2004; Richards et al., 2015) and, occasionally, mortalities in Eastern oysters cultured in U.S. East coast hatcheries (Kehlet-Delgado et al., 2017). In addition to Pacific and Eastern oysters, this pathogen affects commercially important Kumamoto oysters (*C. sikamea*), greenshell mussels (*Perna canaliculus*), and geoduck clams (*Panopea generosa*) (Elston et al., 2008; Estes et al., 2004; Kesarcodi-Watson et al., 2009; Richards et al., 2015). Furthermore, pathogenic *V. coralliilyticus* has been identified globally as a deadly pathogen in various marine species, including finfish, corals, and several bivalve species (Alves Jr et al., 2009; Austin et al., 2005; Jeffries, 1982; Kim et al., 2020).

Antibiotic interventions against bacterial infections in aquaculture are either not available or restricted due to the risks of promoting widespread anti-microbial resistance in bacterial populations (Cabello et al., 2013; Kesarcodi-Watson et al., 2008); furthermore, prophylactic treatments, such as vaccines and probiotics, are either not feasible or not yet commercially available for molluscan aquaculture (Pérez-Sánchez et al., 2018). This is in contrast to the many probiotic products used in crustacean and finfish aquaculture (reviewed by El-Saadony et al., 2021).

The definition of probiotics has been expanded in aquatic environments to reflect that they are not solely antagonists of pathogens, but also can interact positively with both the environment and their hosts. Verschuere et al. (2000), for example, state ”[…] a probiotic is defined as a live microbial adjunct which has a beneficial effect on the host by modifying the host-associated or ambient microbial community, by ensuring improved use of the feed or enhancing its nutritional value, by enhancing the host response towards disease, or by improving the quality of its ambient environment”.

Included in this definition are probiotic-induced modifications of the microbial community that affect host life stages known to be influenced by microbial cues. Bivalves undergo metamorphosis and settlement, where they permanently transition from planktonic larvae to sessile juveniles referred to as spat. Metamorphosis often results in high mortality (Durland et al., 2019). Some biofilms (Campbell et al., 2011; Devakie & Ali, 2002; Rodriguez-Perez et al., 2019; Tritar et al., 1992; Wieczorek & Todd, 1998; Zhao et al., 2003), individually added bacterial isolates (Freckelton et al., 2017), and bacterial supernatants (W. K. Fitt et al., 1990; William K. Fitt et al., 1989; Walch et al., 1999) improve larval oyster metamorphosis, while others result in inhibitory effects (Devakie & Ali, 2002; Dobretsov et al., 2006). Therefore, studies on the use of probiotics in bivalve aquaculture should include their effects on larval settlement and metamorphosis.

This study aimed to develop a combination of beneficial probiotic bacterial isolates that reduced acute mortalities of early larvae of Pacific oysters (*Crassostrea gigas*) resulting from exposure to pathogenic *V. coralliilyticus* strain RE22. We further evaluated whether single or repeated probiotic additions resulted in longer-term benefits to larval growth and metamorphosis of Pacific oysters. Lastly, we determined the effect of a single dose of the combined probiotics on metamorphosis of the Miyagi and Midori strains of the Pacific oyster as well as the Kumamoto oyster (*Crassostrea sikamea*).

## 2. Methods

### 2.1. Isolation and initial screening of probiotic candidates

All experiments were conducted at Oregon State University’s facilities, including the research hatchery at the Hatfield Marine Science Center (HMSC) in Newport, Oregon, USA. Over 311 bacterial isolates were collected from water samples, microalgae tanks, oyster feces, mantle, gills, the gastrointestinal tracts of healthy adult bloodstock and cultures of juvenile spat and larvae of Pacific oysters (*C. gigas*) from the HMSC research hatchery. Additional samples also originated from a commercial hatchery in Netarts Bay, Oregon, and an oyster farm in Yaquina Bay, Oregon. The final probiotic candidates originated from the following samples: *C. gigas* spat that survived a naturally occurring mortality event (D16 and DM14) and gastrointestinal swabs from adult *C. gigas* (B1 and B11). Isolates were enriched in culture broth before being streaked on agar plates of modified, seawater-based Luria-Bertani agar (LBSw; 10 g of tryptone, 5 g of yeast extract, and 15 g agar per liter of filtered seawater) and incubated at 25 °C for at least 24 hours. Plated colonies selected for further screening were grown in LBSw broth (10 g of tryptone, 5 g of yeast extract per liter of filtered seawater) and agitated at 25 °C and 40 RPM on a roller drum (New Brunswick TC-7; New Brunswick Scientific, USA) for 16 to 24 hours. Isolates were cryopreserved in 15% glycerol and stored at -80 °C until further use. All seawater used for bacterial culture was filtered (1 µm, Pentair, Golden Valley, MN) and autoclaved.

Isolates were first screened for their ability to inhibit the growth of *V. coralliilyticus* strains RE22 (Richards et al., 2021) and RE98 (Richards et al., 2014) on LBSw agar plates. Freezer stocks were re-streaked onto LBSw-agar plates and incubated at 25 °C for 24 hours. One colony of each candidate was inoculated into LBSw broth and grown with agitation for at least 24 hours. Five µl of the overnight candidate culture were then dropped onto fresh lawns of approximately 10^4^ CFU of *V. coralliilyticus* RE22 and/or RE98. These 10 cm-diameter LBSw-agar plates were incubated overnight at 25 °C before being checked for zones of inhibition between the probiotic candidates and *V. coralliilyticus*. In addition, putative probiotic candidates were screened on *Vibrio*-selective Thiosulfate Citrate Bile Salt Sucrose (TCBS; Becton Dickinson or Sigma-Aldrich, USA) agar to identify and exclude *Vibrio* spp. during this initial screening process (project workflow in S1 Figure).

### 2.2. Culture of probiotic bacterial isolates and the pathogen *Vibrio coralliilyticus* strain RE22 for use in oyster larval assays

Fresh cultures of bacteria were prepared from cryopreserved samples for every oyster larval experiment. *V. coralliilyticus* strain RE22 was used for all experiments in this study. This strain was isolated from a commercial shellfish hatchery in the Pacific Northwest and is highly pathogenic, often resulting in larval mortalities of 100% at 26 °C (Estes et al., 2004). All probiotic candidates that remained in the evaluation after the initial screening were grown as described above. The bacteria were then washed twice via centrifugation at 3900 x g, resuspended in autoclaved seawater, and the optical density (OD) measured at 600 nm with a spectrometer (Beckman DU 530). With the assumption that OD 1.0 refers to approximately 8 x 10^8^ CFU/ml for all strains used, the cultures were then diluted with autoclaved seawater to a final concentration in the larval culture water of 6 x 10^3^ CFU/mL for *V. coralliilyticus* and 1 x 10^4^ CFU/mL for each probiotic candidate unless otherwise specified.

### 2.3. Production of Pacific oyster (*C. gigas*) and Kumamoto oyster (*C. sikamea*) D-larvae

Oyster broodstock was conditioned and strip-spawned to collect gametes (Langdon et al., 2003). The eggs were fertilized and incubated at a density of 50 to 100 eggs/mL in 30 L containers filled with seawater at 25 °C, 32 ± 2 ppt salinity, and pH of 8.2 ± 0.1. All seawater was pumped from the Yaquina Bay, Newport, passed through 10 µm bag filters, and aerated with soda lime air overnight to adjust the pH. In the experiments described in 2.4. and 2.5., the larvae were hatched in autoclaved seawater with 2 µg/mL chloramphenicol and 10 µg/mL ampicillin. For the experiment described in 2.7, gametes were collected aseptically using ethanol for gonad surface-disinfection and sterile instruments (Douillet & Langdon, 1993), and the incubation seawater was autoclaved.

After incubation, one-day-old (24 hours post-fertilization; hpf) D-larvae were collected on a 45 µm sieve and thoroughly rinsed with sterile seawater. Larvae from different parental crosses were pooled in equal proportions and then distributed among culture containers at a density of 5 to 35 larvae per mL. The culture containers varied from 1 mL to 30 L depending on the type of assay. Twenty-four hpf D-larvae were used in all the infection assays.

### 2.4. Screening putative probiotic candidates for potential pathogenicity against *C. gigas* (Miyagi stock) larvae using well-plate assays

Probiotics were tested for pathogenicity as described by Estes et al. (2004), with minor modifications. A suspension of 24 hpf D-larvae was diluted to 35 larvae per mL with 10 µm-filtered and autoclaved seawater, and one mL of the larval suspension was added to each well of 24-well plates (Corning, USA). Candidate probiotics were each added at a concentration of 3 x 10^4^ CFU/mL. Each treatment was replicated six times. After 48 hours of incubation at 25 °C, larvae were preserved in their wells with 0.1% (v/v) buffered formalin (pH 8.2). Live and dead larvae were counted by light microscopy (Leica DMIL LED inverted microscope, x10 objective). Tissues of dead larvae degraded rapidly, facilitating differentiation between live and dead larvae. If greater than 90% of tissue remained within the shells, the larvae were classified as having been alive. If less than 90% of tissue remained, the larvae were classified as having been dead before formalin-preservation (Madison et al. 2022).

Each assay in this study contained a larvae-only negative control, which did not receive any bacteria and was used to normalize larval mortalities that were unrelated to experimental treatments. This control enabled calculation of relative percent survival (RPS). RPS was calculated as RPS = [1-(percent mortality of treatment group / percent mortality of untreated control group)] x 100. In addition, positive controls were included with larvae that were exposed to *V. coralliilyticus* without addition of probiotics.

### 2.5. Screening putative probiotic candidates for protection of *C. gigas* (Miyagi stock) larvae against *V. coralliilyticus* strain RE22 using well-plate assays

Protective activities of the probiotic candidates were determined in well-plate assays by adding the probiotic candidates (3 x 10^4^ CFU/mL final probiotic concentration) to 24 hpf D-larvae in sterile seawater, followed at 48 hpf by addition of *V. coralliilyticus* strain RE22 at a concentration of 6 x 10^3^ CFU/mL. The larvae were then incubated at 25 °C for 48 hours, after which time they were preserved with 0.1% buffered formalin. Live and dead preserved larvae were observed and counted by light microscopy (Leica DMIL LED inverted microscope, x10 objective).

### 2.6. Identification of probiotic candidates by 16S rRNA gene sequencing

Genetic identification of the candidate probiotics D16, DM14, B1, and B11 was conducted by 16S RNA gene sequencing and the NCBI’s BLAST suite (Altschul et al., 1990). DNA was extracted using phenol:chloroform extraction from one mL of overnight culture according to published protocols with the slight modification that after the final thaw during the RNAse step, 20 µg/mL RNAse A was added, and samples were incubated at 34°C for 30 min (Crump et al., 2003). Amplification of the 16S rRNA gene from each genome was performed under standard PCR conditions with the forward primer 8F 5’-AGAGTTTGATCCTGGCTCAG and the reverse primer 1513R 5’-ACGGCTACCTTGTTACGACTT amplifying an approximately 1500 bp piece of DNA. Dideoxy sequence reads were generated from the cleaned PCR product using the same primers. The forward and reverse sequence reads were assembled and trimmed, and the resulting consensus sequence was then queried against the NCBI’s 16S ribosomal RNA sequence database using BLASTN (Altschul, 1990).

### 2.7. Developing a probiotic combination treatment using well-plate assays

Promising candidates were combined for testing to take advantage of potentially several concurrent, probiotic modes of action. Each probiotic test well was filled with a one mL suspension of approximately 35 one-day-old (24 hpf) D-larvae and probiotic isolates (B11, DM14, and D16) in sterile seawater. These probiotics were tested individually or combined in equal concentrations of two or three probiotics to achieve final total probiotic concentrations of 3 x 10^4^ CFU/mL. At 48 hpf, the probiotic-treated larvae were challenged with *V. coralliilyticus* strain RE22 and incubated until 96 hpf. A positive control consisted of larvae with additions of *V. coralliilyticus* alone (Vcor only). Lastly, a negative control was included that consisted of larvae that did not receive probiotics nor *V. coralliilyticus* (Larvae only). At 96 hpf, larvae were preserved with the addition of buffered formalin and live and dead larvae were counted, as described above.

### 2.8. Effects of the probiotic combination on size, settlement and metamorphosis of *C. gigas* (Miyagi stock) larvae in a long-term assay

#### 2.8.1. Larval culture and experimental treatments

A new cohort of larvae (24 hpf) from a separate spawn was stocked at a concentration of five larvae per mL in 10-L containers filled with 10 µm-filtered seawater, with five replicates per treatment group. One treatment group received probiotics added to the larval culture water once at 24 hpf (Single PB Addition). The other treatment group received probiotics following water changes every 48 hours (Multiple PB Additions). A control treatment of larvae alone (Larvae Only), without additions of probiotics, was also included.

Probiotics were prepared as previously described but were applied to the 10-L cultures at a concentration of 6 x 10^4^ CFU/mL each, resulting in a total combination treatment of 1.8 x 10^5^ CFU/mL. This higher probiotic concentration was necessary because assays conducted in non-sterile, 10 µm-filtered water often resulted in reduced beneficial probiotic effects (unpublished data). Coarsely filtered seawater was chosen for these assays as it is typically used for rearing larvae in commercial shellfish hatcheries.

The first water change occurred 72 hours after stocking the larvae, then every 48 hours after that. During water changes throughout this experiment, larvae were poured over a 45 µm-screen to avoid removing slow-growing larvae. Larvae were rinsed with 10 µm-filtered seawater at each water change. As part of the water change, the containers were scrubbed with 0.02% (v/v) Vortexx (Ecolab, USA) and thoroughly rinsed with hot tap water. In addition, the airlines were rinsed with tap water before the containers were refilled with fresh seawater. Metamorphosis success was determined on 20, 22 and 24 dpf.

Larvae were cultured according to established methods (Langdon et al., 2003). Briefly, larvae were fed during the first six days on an algal diet of the flagellate *Tisochrysis lutea* (strain T-ISO) at 4 x 10^4^ cells/mL. On day seven, the diet was modified to include a 50/50 mixture (by cell concentration) of T-ISO and the diatom *Chaetoceros gracile* and the total cell concentration was increased to 5 x 10^4^ cells/mL and then further increased to 8 x 10^4^ cells/mL on day twelve.

#### 2.8.2. Larval size

Random larval sub-samples were collected from the 10-L containers on day 8 and transferred to 24-well plates in an attempt to carry out a challenge assay with larvae exposed to *V. coralliilyticus* over a four-day period. This assay failed because no mortalities were observed in any treatments or the positive control (Vcor only) (data not presented). Control larvae from wells that were not exposed to *V. coralliilyticus* RE22 were sampled on day 12, preserved with 0.1% phosphate-buffered formalin (pH 8.2), and photographed at 40X objective magnification (Leica DMIL LED inverted microscope; Leica DFC400 camera; Leica Application Suite 4.8 Leica, Germany). Over 350 larvae were measured from each treatment consisting of six replicates. Shell lengths (defined as the greatest dimension parallel to the shell hinge) of larvae were measured using the software Image Pro Premier 9 (Media Cybernetics, USA).

#### 2.8.3. Larval settlement and metamorphosis

Sixteen days post-fertilization (dpf), 4 x 4-inch marble tiles were added to each 10-L container, and larvae retained on a 240 µm-screen began setting naturally. On 20 dpf, the marble tiles were removed and photographed. The settled larvae on each tile were counted using the imaging analysis software Image Pro Premier 9 (Media Cybernetics, USA). In addition, larvae set on the containers and airlines were counted manually during each water change. Spat that set on the containers, beakers, and tiles were categorized as naturally set.

After removing the tiles containing the spontaneously settled larvae, 2 x 10^-4^ M epinephrine (Epi) was used to induce the metamorphosis of remaining larvae on days 20, 22, and 24 (Coon et al., 1986). Non-metamorphosing larvae were returned to their culture containers on days 20 and 22. Any larvae that had not metamorphosed after exposure to epinephrine on day 24 were counted as larvae. Metamorphosed larvae were moved to a flow-through upweller system, where they were cultured until 30 dpf to allow production of additional shell growth. These spat were categorized as successfully metamorphosed with epinephrine, preserved in 0.1% phosphate-buffered formalin (pH 8.2), and counted.

### 2.9. Effects of the probiotic combination on larval settlement and metamorphosis of the Miyagi and Midori stocks of the Pacific oyster (*Crassostrea gigas*) and the Kumamoto oyster (*Crassostrea sikamea*) in long-term assays

The experiment described above (section 2.8) was repeated with new cohort of larvae from two different stocks of the Pacific oyster - the commonly farmed Miyagi stock and the newly introduced Midori stock (de Melo et al., 2021), and larvae of the Kumamoto oyster. The experimental conditions were similar to those described above (single PB addition) with a few differences: 1) the larvae were raised in 30 L of seawater, and the first water change occurred 48 hours after stocking the larvae, 2) live *Nannochloropsis occulata* (Nanno) cells were added to the T-ISO diet on days two to six, 3) the algal diet was checked daily and, if the larvae had consumed all food, the total cell concentration of the algal diet was increased by 5 x 10^3^ cells/mL from an initial concentration of 3.5 x 10^4^ cells/mL T-ISO, and 1.5 x 10^4^ cells/mL Nanno and, 4) larval densities were intentionally reduced during the experiment by sieving and discarding slow-growing larvae as per best practices of commercial oyster hatcheries (Barton et al. 2012).

Larvae were initially stocked as described in 2.8. Slow-growing larval *C. gigas* were removed from the cultures if they were not retained on a 64 µm-screen on day 7 or on an 80 µm-screen on day 9. Larval *C. sikamea* are smaller than larval *C. gigas*; therefore, the screening procedure was adjusted accordingly. Larvae were reduced to a density of one larva per ml on day 9. Slow-growing larvae not retained on a 180 µm-screen on day 17 were removed. Generally, larvae grew faster in this experiment, so the tiles were added on day 15 and removed on day 17 post-fertilization, and three rounds of epinephrine were used on days 17, 19, and 21 post-fertilization.

### 2.10. Statistical analyses

Data from larval survival assays was converted to RPS, as described in 2.4. RPS values were arcsine square root transformed prior to analysis. Statistical analyses were conducted using R statistical software (Version 4.0.3, R Project for Statistical Computing). Normality was assessed using the Shapiro-Wilk test and Q-Q plots. Homogeneity of variance was assessed using Levene’s test.

For normally distributed, homoscedastic data sets, multiple comparisons of treatment group means against control groups were conducted using Dunnett’s test. When multiple comparisons between all treatment group means were of interest, data was first fitted to a linear model, then a one-way ANOVA was conducted. If ANOVA showed significant differences (*P* < .05), Tukey’s HSD test was conducted at a .05 level of significance. Comparisons between two treatment group means were conducted using a two-sample t-test. When heteroscedasticity was observed, Welch’s two sample t-test was used.

Nonparametric methods were used when violations of normality and variance assumptions were observed. For multiple comparisons of treatment groups, the Kruskal-Wallis one-way ANOVA was conducted. When significant differences (*P* < .05) were found, Dunn’s test with the Benjamini-Hochberg correction was used to make pairwise comparisons. Comparisons between two treatment groups were conducted using the Mann-Whitney U test.

## 3. Results

### 3.1. Initial screening of probiotic candidates

From all microbial strains collected for probiotic screening, approximately 28.3% were not revivable after being frozen with glycerol or failed to sufficiently grow in LBSw within 48 hours, 36.7% grew on TCBS agar and were consequently excluded, leaving 35% screened on agar plates against *V. coralliilyticus* strain RE22. Ultimately 13 strains were suggestive of contact inhibition or zones of clearing on the *V. coralliilyticus* lawn. These strains proceeded to pathogenicity testing with oyster larvae (project workflow in S1 Figure).

### 3.2. Screening putative probiotic candidates for potential pathogenicity against *C.gigas* (Miyagi stock) larvae using well-plate assays

Putative probiotic candidates DM7, DM36, DM39, and D20 passed the agar inhibition test, but when added to larvae 24 hpf, caused a reduced survival that was statistically lower than that of the non-probiotic control (Larvae Only). As a result, these candidates were excluded from further evaluation (S1 Table).

### 3.3. Screening putative probiotic candidates for protection of *C. gigas* (Miyagi stock) larvae against *V. coralliilyticus* strain RE22 using well-plate assays

The highly virulent *V. coralliilyticus* strain RE22 was added to two-day-old larvae (48 hpf) in well plate assays, at a concentration of 6x10^3^ CFU/mL 24 hours after probiotic additions at 3 x 10^4^ CFU/mL. The positive control (Vcor only) did not receive any probiotics and resulted in almost complete larval mortality, reducing the relative percent survival to an average of 3.40% (Figure 1).

**Figure 1:**
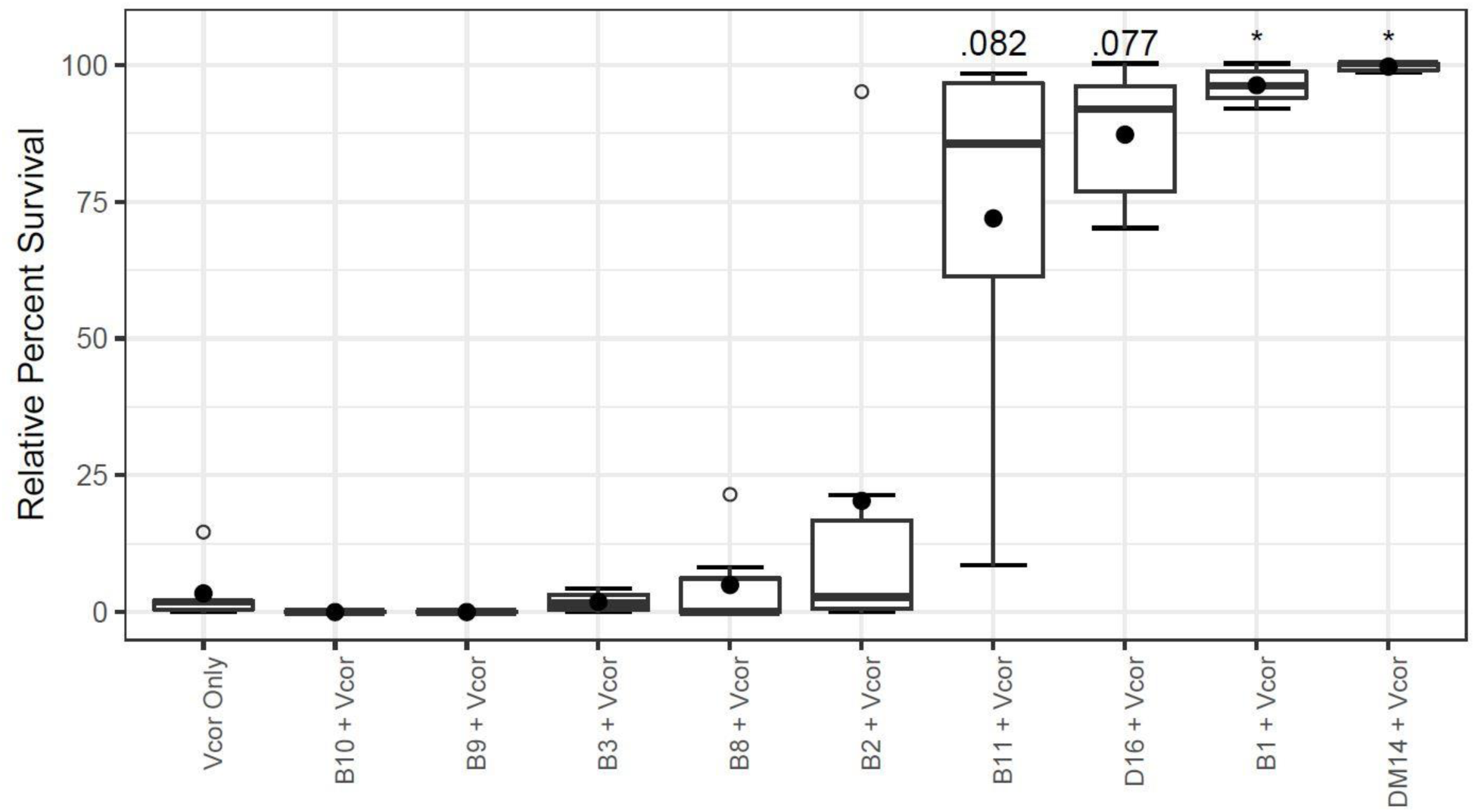
*C. gigas* larvae were protected against *V. coralliilyticus* RE22 (Vcor) with probiotic candidates B11, D16, B1, and DM14. Probiotic candidates were evaluated in well-plate assays with 24 hpf-old *C. gigas* larvae. “Vcor Only” was the positive control with no additions of probiotics but was inoculated with *V. coralliilyticus* 48 hpf. A negative control did not receive any probiotics or pathogen and was used to normalize mortalities that were not related to pathogen or probiotic treatments to calculate relative percent survival (RPS) values. Filled circles depict the average relative percent survival of six replicate wells. The boxes indicate the upper and lower quartiles, and the bar represents the median or middle quartile. The ends of the whiskers represent the most extreme values within the 1.5x interquartile range (IQR), and the empty circles indicate outliers. Only the treatments using the probiotic candidates B11, D16, B1, and DM14 yielded more than 68% improved average RPS, compared to the positive control (Vcor Only). However, statistical analysis using Dunn’s Test with a Benjamini-Hochberg Correction resulted in no (D16 *P*=.077; B11 *P*=.082) or low (B1/DM14 .05*>P*>.01; indicated with asterisks) statistically significant differences between larval survival with the probiotic treatments and the “Vcor Only” control, due to the high variance of some of the treatments (S2 Table).

Of the nine remaining candidates tested in this experiment, additions of the four probiotic candidates DM14, B1, D16, and B11 each resulted more than a 68% increase in mean RPS compared to that with the positive *V. coralliilyticus* control (Vcor Only) (S2 Table). Treatments with DM14 or B1 resulted in minimal larval losses and an average survival of 99.71 ± 0.87% and 96.29 ± 3.30%, respectively (S2 Table). In contrast, larvae treated with D16 or B11 showed high variabilities in survival among replicates (S3 Table). Larvae treated with D16 had a slightly reduced survival in two replicate wells (71.8% and 70%), but high survival in the remaining four test wells leading to a mean RPS of 87.28 ± 12.96% (S2 and S3 Tables). Lastly, larvae treated with B11 had high survival in three wells (95.31%, 96.77%, 98.15%), moderate in two wells (56.4% and 75.47%), and a low survival of 8.5% in one well, leading to a mean RPS of 71.96 ± 35.10% (S2 and S3 Tables). Repeated trials, including the use of different seawater sources (data not shown) did not reduce inter-replicate variabilities observed for all probiotic candidates. Probiotic candidates were, therefore, advanced to further testing when mean RPS values were at least 40%, regardless of whether a statistical significance of *P*<.05 was achieved. The addition of candidates B2, B3, B8, B9, and B10 did not result in an 40% improved mean RPS compared to the positive control (Vcor Only), and these isolates were consequently excluded from further testing (S2 Table).

### 3.4. Identification of the probiotic candidates by 16S rRNA gene sequencing

After the screening of the putative probiotics against *V. coralliilyticus* in larval assays, promising probiotic candidates were submitted to 16S rRNA sequencing. BLASTN suite search identified D16 and DM14 as different *Pseudoalteromonas* spp., and B11 as an *Epibacterium* sp. Finally, isolate B1 was identified as a *Vibrio* sp. and was consequently excluded at this stage. All 16S sequences were deposited to Genbank (Accessions ON863687-ON863692).

### 3.5. Developing a probiotic combination treatment using well-plate assays

Individual or combination treatments of the candidate probiotics B11, D16, and DM14 were evaluated for their ability to reduce larval mortalities due to *V. coralliilyticus* and to decrease variability among replicates (Figure 2).

**Figure 2:**
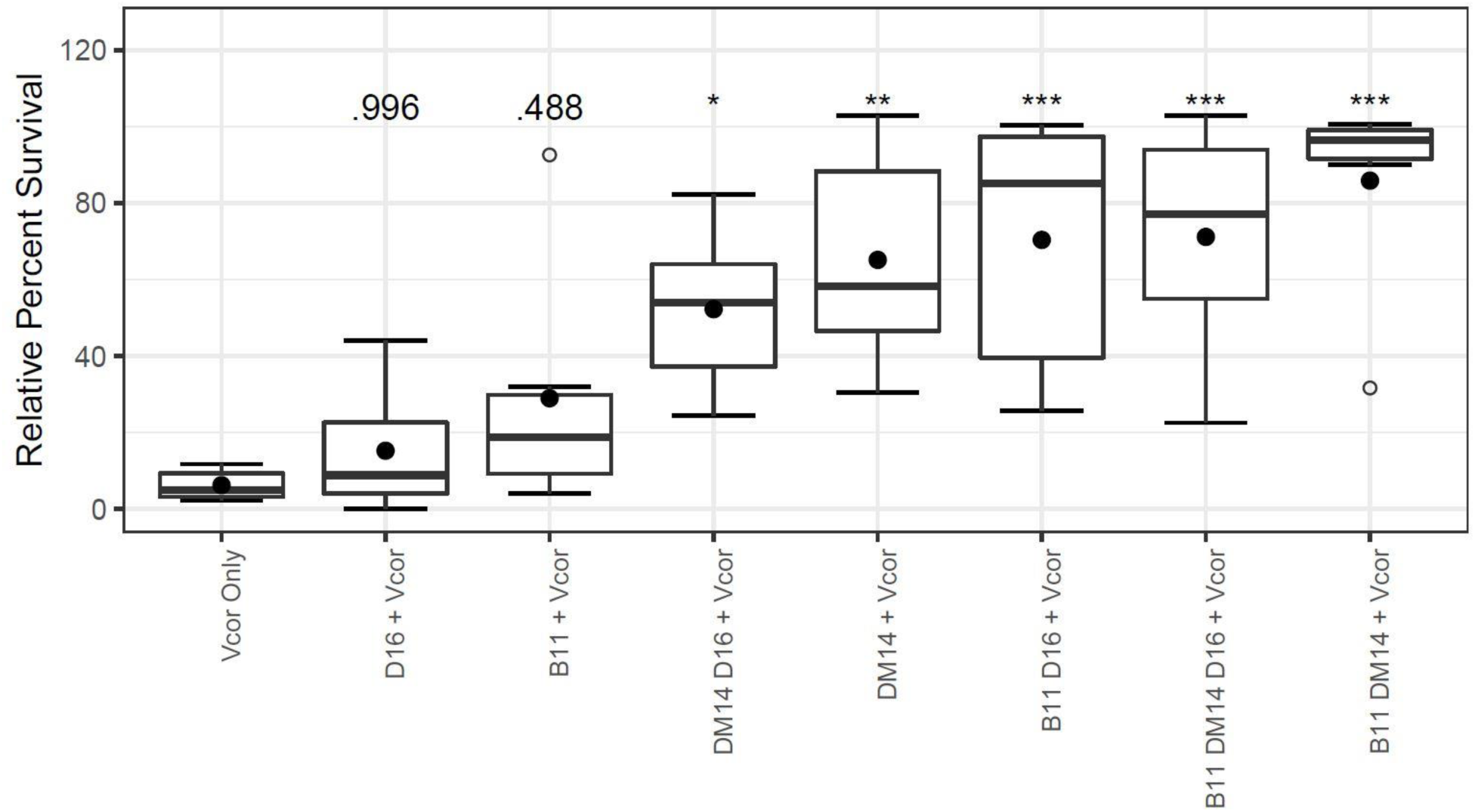
Relative percent larval survival with individual or combination probiotic treatments against *V. coralliilyticus* strain RE22 for 1-day-old D-larvae of *C. gigas*. Individual or combinations of probiotics were evaluated for their ability to decrease larval mortalities due to *V. coralliilyticus*. “Vcor Only” was the positive control that did not receive any probiotics. Filled circles depict the average relative percent survival of six replicate wells. The boxes indicate the upper and lower quartiles, and the bar represents the median or middle quartile. The ends of the whiskers represent the most extreme values within the 1.5x interquartile range (IQR), and the empty circles indicate outliers. *, ** and *** indicate statistical differences from “Vcor Only” at *P*≤.05, *P*≤.01, and at *P*≤.001, respectively.

Oyster larvae, that did not receive any probiotics 24 hpf but were challenged with *V. coralliilyticus* 48 hpf, showed a low relative percent survival (RPS) averaging 6.23% across the six replicate wells (S5 Table). Individual applications of D16 and B11 did not increase the mean RPS of the larvae compared to the positive control (Vcor only) (15.24%; *P*=.996) and (28.98%; *P*=.488), respectively. Only the individual probiotic candidate DM14 significantly increased the mean RPS (65.17%; *P*=.003). However, when D16 and B11 were combined the mean RPS was 64.17% higher than the positive control (*P*=.001), and combining DM14 and D16 resulted in a 46.05% improvement (*P*=.034). Combining B11 and DM14 achieved a mean RPS of 85.88% (*P*<.001); however, one of the replicates was categorized as an outlier (RPS of 31.67%; data not shown). The three-strain combination resulted in a mean RPS of 71.18% (*P*=.001) (S5 Table). Overall when the results of the individual, two-strain, and three-strain combinations were combined (S2 Figure), there was no statistical significance of the single strain additions compared to the positive control (*P*=.085). The effects of the two- and the three-strain combinations were similar to each other (*P*=.999) but significantly different from that of the Vcor only control (S6 Table).

### 3.6. Effects of the probiotic combination on size, settlement and metamorphosis of *C.gigas* (Miyagi stock) larvae in long-term assays

#### 3.6.1. Larval size

Larvae from the five replicate control containers that did not receive any probiotic treatment had an average shell size of 153.21 ± 28.52 µm, which was significantly smaller than those of the treatment groups that received either a single dose or multiple doses of probiotics (Dunnett’s Test, *P*≤.001) (Table 1). The treatment group that received repetitive probiotic doses after each water change had an average shell size of 161.24 ± 30.89 µm, and the treatment group that received a single probiotic addition 24 hpf resulted in a value statistically similar to the repetitive dosing treatment (*P*=.258), but with slightly greater average shell length of 164.49 ± 30.81 µm (Table 1) (S3 Figure).

**Table 1:** Average shell lengths of *C. gigas* (Miyagi stock) larvae after probiotic treatments.

| Dunnett's Test (comparing to Larvae Only control) |  |  |  |  |  |
| --- | --- | --- | --- | --- | --- |
| Treatment | Mean shell Length (μm) | Standard Deviation | Difference | 95% Confidence Interval (CI) | p-value |
| Single PB Addition | 164.49 | 30.81 | 11.28 | (6.57, 16.00) | <.001*** |
| Multiple PB Additions | 161.24 | 30.89 | 8.03 | (3.15, 12.92) | .001*** |
| Larvae Only | 153.21 | 28.52 |  |  |  |
| Tukey-Kramer (pairwise comparisons between PB treatments) |  |  |  |  |  |
|  | Pairwise Comparison |  | Difference | 95% CI | p-Value |
|  | PB Multiple - PB Single |  | -3.25 | (-8.09, 1.60) | .258 |

\*\*\* indicates $P \leq .001$ ; probiotic (PB) = combination of B11, D16, and DM14

#### 3.6.2. Larval settlement and metamorphosis

Spat were categorized as undergoing successful metamorphosis when they survived until 30 dpf, i.e., for at least six days after settlement or addition of epinephrine. An average of 2.24% of larvae in the control treatment group that did not receive any probiotics naturally set on the marble tiles and surfaces of the culture container (Figure 3) (S7 Table). The remaining larvae exposed to epinephrine displayed a successful metamorphosis rate of 4.81%, resulting in a combined metamorphosis rate of 7.06%. A one-time probiotic addition at 24 hpf resulted in 8.29% natural settlement and metamorphosis. When epinephrine was added to the remaining larvae, 13.03% metamorphosed, resulting in a combined metamorphosis rate of 21.32% that was significantly higher than that of the control without addition of probiotics (*P*=.004). Repeated dosing of the probiotic combination after each water change resulted in a successful settlement and metamorphosis rate of 2.8% of larvae on marble tiles. The addition of epinephrine yielded a metamorphosis rate of 4.24%, resulting in a total metamorphosis rate of 7.04%, which was not significantly different from the “Larvae Only” control (*P*>.999) (S7 Table).

**Figure 3:**
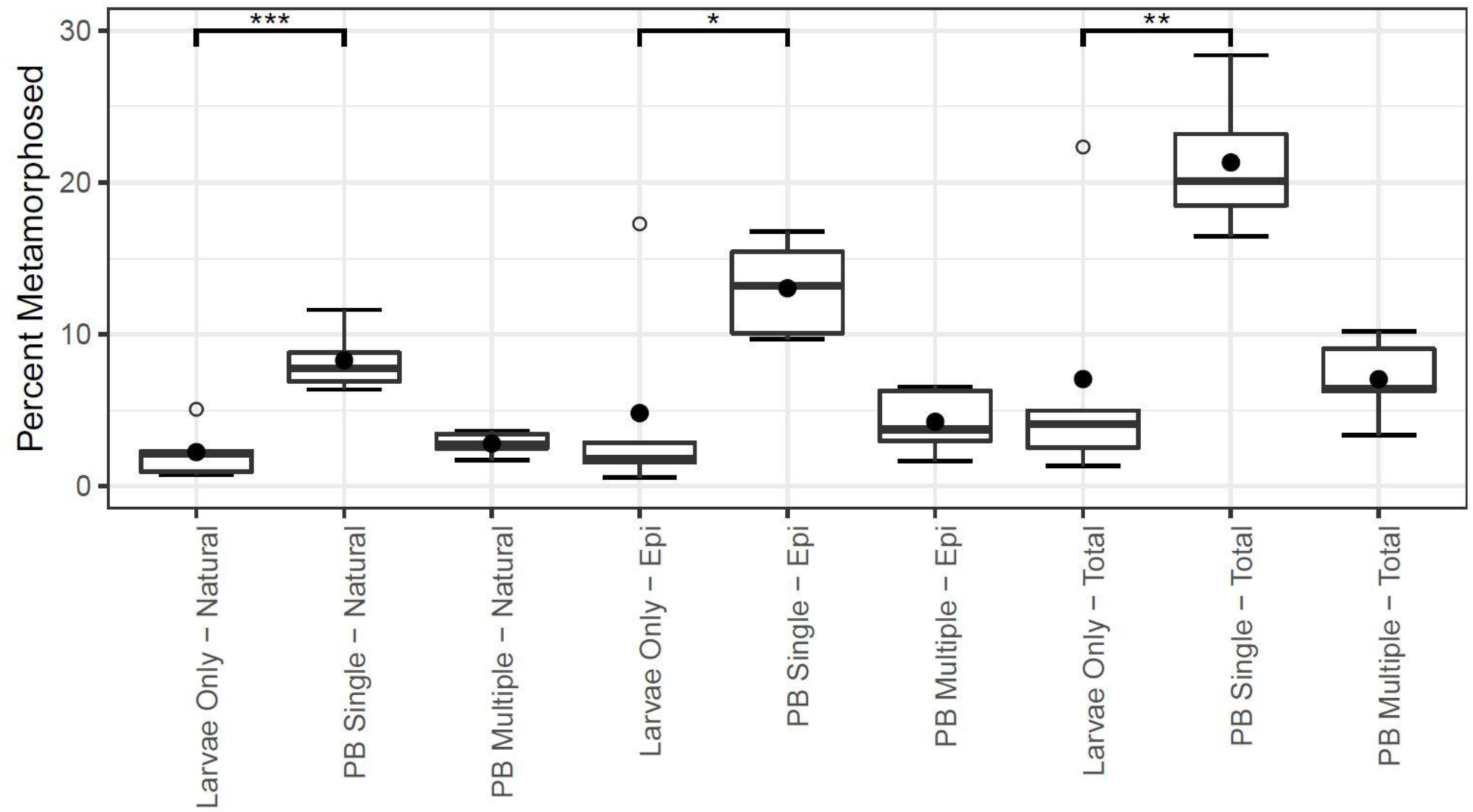
Single application of the probiotic (PB Single) improved settlement and metamorphosis of *C.gigas* (Miyagi stock) larvae. A single dose of the probiotic combination (PB Single) 24 hpf significantly improved successful metamorphosis of naturally set (*P*<.001***) and epinephrine-treated (*P*=.028*) *C. gigas* larvae, resulting in significantly improved total metamorphosis (*P*=.004**), compared with the control with no probiotic additions. Metamorphosis success with repeated probiotic applications (PB Multiple) did not differ from that of control larvae (Larvae Only) (*P*>.999) (S7 Table). Filled circles depict the average relative percent survival of five replicate 10-L containers. The boxes indicate the upper and lower quartiles, and the bar represents the median or middle quartile. The ends of the whiskers represent the most extreme values within the 1.5x interquartile range (IQR), and the empty circles indicate outliers.

### 3.7. Effects of the probiotic combination on larval settlement and metamorphosis of *C. gigas* (Miyagi and Midori stocks) and *Crassostrea sikamea* (Kumamoto) in long-term assays

The addition of a single dose of the B11, DM14, D16 probiotic combination 24 hpf increased the percentage of larvae that metamorphosed successfully in Miyagi and Midori stocks of *C. gigas* as well as the Kumamoto oyster (*P*≤.02) (Table 2). Because the larval density was reduced to one larva per mL on day nine post-fertilization, the percent metamorphosed was based on the number of larvae restocked on that day.

**Table 2:** Settlement and metamorphosis of larvae from two stocks of *C. gigas* (Miyagi and Midori) and the Kumamoto oyster (*C. sikamea*) were improved with a single addition of the probiotic combination B11, D16, and DM14 at 24 hpf.

| Oyster species/stocks | Treatment | Natural Set Avg. % (±%STDEV) | Metamorphosis with Epi Avg. % (±%STDEV) | Total Metamorphosed Avg. % (±%STDEV) | Two-sample t-test $p$-value<sup>b</sup> |
| --- | --- | --- | --- | --- | --- |
| Miyagi<sup>a</sup> | w/ probiotics | 24.02 (±11.18) | 55.74 (±10.51) | 79.76 (±6.03) | .020 |
| Miyagi<sup>a</sup> | no probiotics | 20.89 (±6.18) | 43.21 (±8.38) | 64.10 (±10.41) |  |
| Midori | w/ probiotics | 22.21 (±11.82) | 34.46 (±6.54) | 56.67 (±6.37) | .015 |
| Midori | no probiotics | 15.34 (±2.44) | 29.36 (±7.42) | 44.70 (±5.81) |  |
| Kumamoto | w/ probiotics | 26.18 (±3.79) | 31.72 (±4.99) | 57.90 (±4.04) | .004 |
| Kumamoto | no probiotics | 21.31 (±2.82) | 24.82 (±4.20) | 46.14 (±5.03) |  |

<sup>a</sup>Miyagi is the stock name of the commonly cultured Pacific oyster (*C. gigas*) on the US West coast, while the Midori stock was introduced to the US West coast in the early 2000s (de Melo et al., 2021); <sup>b</sup>Two-sample t-tests were conducted on the total ratios of set larvae. All other t-tests in S8 Table.

#### 3.7.1. Effects on *C. gigas* Miyagi stocks

Larvae in the control treatment group that did not receive any probiotics showed a rate of 20.89% natural settlement and metamorphosis on marble tiles (S4 Figure). Non-settled larvae were then treated with epinephrine and 43.21% underwent successful metamorphosis. The subsequent total metamorphosis of larvae without any probiotics was 64.1%. A one-time probiotic addition 24 hpf resulted in 24.02% natural settlement. When epinephrine was added to the remaining larvae, 55.74% metamorphosed, resulting in a combined metamorphosis rate of 79.76% that was significantly higher than total metamorphosis without addition of probiotics (*P*=.020) (Table 2).

#### 3.7.2. Effects on *C. gigas* Midori stocks

The probiotic treatment resulted in an average settlement and metamorphosis rate of 22.21% for naturally set Midori larvae versus 15.34% for non-probiotic treated controls (S5 Figure). With epinephrine, 34.46% of the larvae successfully metamorphosed compared to 29.36% without a probiotic addition. The total metamorphosis of Midori larvae treated with probiotics was 56.67% versus 44.7% (*P*=.015) (Table 2).

#### 3.7.3. Effects on *C. sikamea* (Kumamoto) oysters

With Kumamoto oysters, the probiotic treatment increased the proportion of successfully naturally set and metamorphosed larvae to 26.18% compared to 21.31% of non-probiotic treated larvae (S6 Figure). The epinephrine treatment resulted in 31.72% metamorphosis versus 24.82% for non-probiotic treated larvae. The total proportion of successfully metamorphosed spat was 57.9% with the addition of probiotics, while non-probiotic treated Kumamoto larvae had a significantly lower total metamorphosis success of 46.14% (*P*=.004) (Table 2).

## 4. Discussion

This study describes the development of a promising novel probiotic treatment for both disease prevention and enhancement of larval development. The best treatment consisted of a combination of three individual beneficial bacterial isolates added to the larvae culture once at 24 hpf. This treatment significantly improved the survival of Pacific oyster larvae exposed to a lethal dose of a highly virulent *V. coralliilyticus* strain RE22 added 24 hours after the probiotic addition. In addition, a single dose of the probiotics 24 hpf resulted in higher larval settlement and metamorphosis success for two stocks of Pacific oysters and for the Kumamoto oyster. Besides mass mortalities during critical stages of larval development, larval metamorphosis and post-settlement survival are common seed production problems for commercial hatcheries (Barton et al. 2012).

Initial selective steps in choosing probiotic candidates in this study included survival of freeze-thaw steps, absence of pathogenicity towards larvae, and the ability to suppress the growth of *V. coralliilyticus* strain RE22 using agar plate assays. In addition, growth on the *Vibrio*-selective agar (TCBS) led to the exclusion of candidates with antagonistic properties. Due to likely intense interspecies competition in this genus (see, for example, Borgeaud et al. 2015), *Vibrio* spp. often displayed moderate to strong inhibition of *V. coralliilyticus* on agar plates and *Vibrio* isolate B1 showed promise in survival assays with oyster larvae. For unknown reasons, B1 failed to grow on TCBS plates at the screening stage and was only subsequently identified as a *Vibrio* sp. by sequencing, leading to its exclusion from further testing. Members of the genus *Vibrio* were intentionally excluded from the probiotic combination as they have the ability to quickly adapt to new environments and acquire virulence factors by horizontal gene transfer that could potentially transform the isolate from an oyster commensal to a pathogen (Bruto et al., 2017; Le Roux & Blokesch, 2018).

Individual probiotic candidates that improved mean RPS in initial screenings by at least 40% compared to a non-probiotic treated control, were selected for inclusion in the probiotic combination. Ultimately, a combination of the three isolates B11, DM14, and D16 significantly increased the efficacy of the treatment. High levels of variation among replicates were observed in these assays irrespective of probiotic isolate, larval rearing method, including larval density and rearing volume (well-plates or 10-L containers), or seawater quality (autoclaved or 10 µm-filtered). Such variation seems common in trials with oyster probiotics (Sohn et al., 2016) and may be due to small, uncontrolled differences in environmental conditions among replicates that affect the bacterial composition of the cultures, microbiome or host-microbe interactions (Stevick et al., 2019).

Application of probiotics in aquaculture systems, including probiotics tested in oyster hatcheries, are commonly dependent on repeated probiotic dosing (Kapareiko et al., 2011; Sohn et al., 2016). It has been suggested that repeated additions are necessary due to the inability of the probiotic bacteria to permanently colonize the animal’s mucosal surfaces, including the gut (Gatesoupe, 1999). This study evaluated whether repeated dosing after each water change was required for prolonged beneficial effects of the probiotic combination. We found larvae to be larger on day 12 when probiotics were added versus larvae from a control with no probiotic additions; furthermore, there were significant improvements in settlement and metamorphosis success for *C. gigas* larvae with a single dose of the probiotic combination at 24 hpf, whereas repetitive dosing was less beneficial. Significantly improved metamorphosis and settlement with a single dose of probiotics at 24 hpf was consistent for larvae from two stocks of *C. gigas* (Midori and Myagi) and *C. sikamea* (Kumamoto) oysters. It is known that bacteria affect bivalve settlement via biofilm formation or secretion of chemical settlement cues (Fitt et al., 1990; Weiner et al., 1985). However, after multiple water changes, it is unlikely that the initial probiotic dosing between 24 and 72 hpf influenced subsequent biofilm formation or settlement cues more than two weeks later.

In the past decade, research efforts have focused on identifying new probiotics intended to increase the survival of oyster larvae exposed to pathogens, e.g., Karim et al. (2013), Kesarcodi-Watson et al. (2012), and Lim et al. (2011). It has been found that the early establishment of the microbiome in oyster larvae is beneficial to early oyster development and survival (Gomez-Gil et al., 2000; Harris, 1993; Kesarcodi-Watson et al., 2012; Prado et al., 2010; Schulze et al., 2006). Knowledge of the beneficial modes of action of probiotics is sparse, particularly in the aquatic environment when probiotic benefits could be related to interactions with the rearing environment, the host-associated microbial community, or with the host’s eukaryotic cells (Gomez-Gil et al., 2000; Lebeer et al., 2010; Prado et al., 2010; Van Doan et al., 2020). These modes could include modulation of the resident microbiome to a more beneficial state, direct antagonism against individual cells of a pathogen, such as killing by direct contact and toxin secretion, production of antimicrobials, or by changing environmental conditions in ways that limit pathogen growth via competitive exclusion (Bhogoju & Nahashon, 2022; Kaushik et al., 2022; Kesarcodi-Watson et al., 2008; Sohn et al, 2016). Probiotics may also directly interact with the eukaryotic cells of the host and contribute to digestion by supplying enzymes, acting as a nutritional source, restoring mucosal integrity and barrier functions, or modulating the immune response (Bhogoju & Nahashon, 2022; Kesarcodi-Watson et al., 2008; Sohn et al., 2016).

It is conceivable that multiple modes of action are present in an individual probiotic or that a combination treatment, as studied in this work, could take advantage of different modes of action. Further advantages of combination treatments could be due to a broader spectrum of activity against different pathogen strains or species, retention of beneficial activity even if a single probiotic constituent fails, and potentially reducing selective pressures against probiotic isolates that decrease benefits.

Further investigations are needed to explore the modes of action of new probiotic treatments in oyster larvae, such as the effect of probiotic additions on the host’s microbiome and the microbial communities of the rearing environment (e.g., Modak & Gomez-Chiarri, 2020; Sohn et al., 2016; Stevick et al., 2019). Such research efforts will help support a sustainable oyster industry facing many challenges due to global warming and ocean acidification.

## Acknowledgments

The authors thank the MBP staff for their support in rearing larvae, Jennifer Hesser for assisting in some of the experiments, MK English for the 16S rRNA gene sequencing of the probiotic candidates. Whiskey Creek hatchery and Oregon Oyster Farms, Oregon, kindly provided biological materials for isolation of candidate probiotics. This work was supported by a NOAA National Sea Grant [NA18OAR4170346] awarded to CS, CL, and RM, and a NOAA Saltonstall-Kennedy grant [NA18NMF4270220] awarded to CL.

**Appendix A. Supplementary figures S1 – S6.**

**Appendix B. Supplementary tables S1 – S8.**

