## Appendix Supplemental Figures for "A marine probiotic treatment against the bacterial pathogen *Vibrio coralliilyticus* to improve the performance of Pacific (*Crassostrea gigas*) and Kumamoto (*C. sikamea*) oyster larvae"

David Madison[a1], Carla Schubiger[b], Spencer Lunda[b], Ryan S. Mueller[c], Chris Langdon[a]

[a]Coastal Oregon Marine Experiment Station and Department of Fisheries, Wildlife, and Conservation Sciences, College of Agricultural Sciences, Oregon State University, Corvallis, Oregon, 97330, USA

[b]Department of Biomedical Sciences, Carlson College of Veterinary Medicine, Oregon State University, Corvallis, Oregon, 97330, USA

[c]Department of Microbiology, College of Science, Oregon State University, Corvallis, Oregon, 97330, USA

[1] Current address: JL. Trans Balauring, Desa Merdeka, Kecamatan Lebatukan, Kab. Lembata, 86681 NTT, Indonesia

### Appendix A. Supplemental figures.

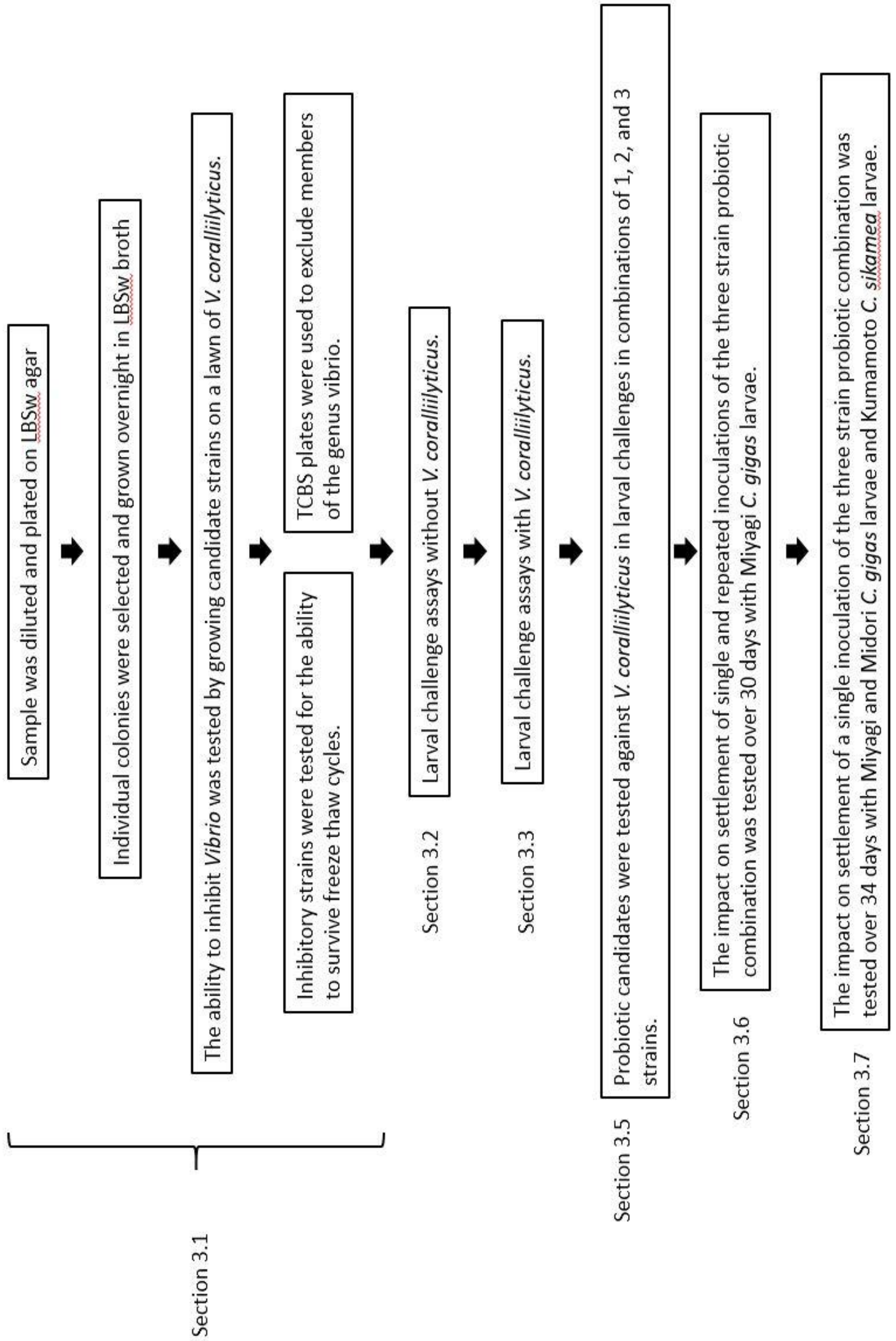


**S1 Figure: Project workflow.**

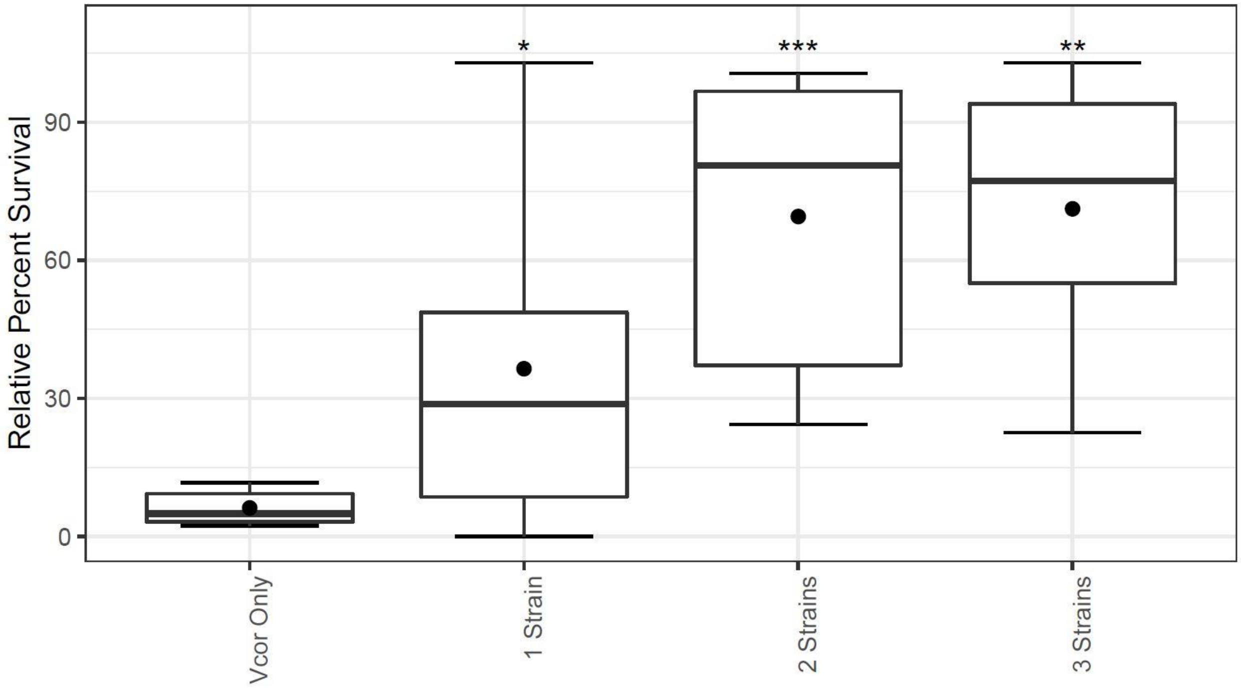


**S2 Figure: Relative percent survival (RPS) of collated individual or combination treatments against *V. coralliilyticus* for 1-day-old D-larvae of *C. gigas*.** Individual or combinations of probiotics were evaluated for their ability to reduce mortalities of 1-day-old *C. gigas* larvae exposed to *V. coralliilyticus* RE22. Data were collated for three individual strains, three combinations of two probiotics (B11/DM14, B11/D16, DM14/D16), and one treatment of B11, DM14, and D16. "Vcor Only" was the no-probiotic positive control. The combinations of the individual strains compared with the positive control showed more than 30% higher RPS (36.43 ± 33.43% STDEV), which was statistically different (*P*=.038*). Combinations of two or all three probiotics resulted in higher RPS values compared to that of the positive control "Vcor only" (2-strain: 69.52 ± 30.15%, *P*<.001***; 3-strain: 71.18 ± 30.72% STDEV, *P*=.002**), but the RPS of 2-strain and 3-strains were not statistically different from each other (*P*=.498) (S6 Table). Filled circles depict the average relative percent survival of six replicate wells. The boxes indicate the upper and lower quartiles, and the bar represents the median or middle quartile. The ends of the whiskers represent the most extreme values within the 1.5x interquartile range (IQR), and the empty circles indicate outliers.

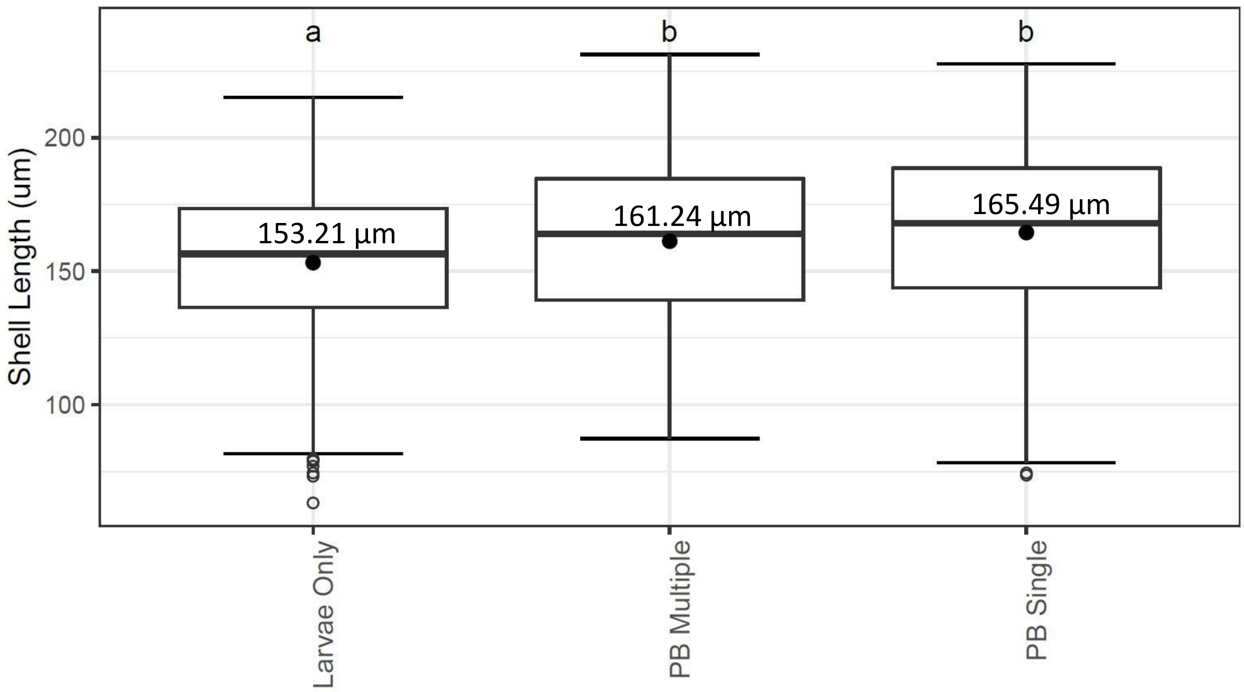


**S3 Figure: Repetitive probiotic dosing is not necessary to significantly increase the shell growth of *C. gigas* larvae.** *C. gigas* larvae were dosed with a single addition of a probiotic combination and their growth compared with that of larvae repetitively dosed every 48 hours after routine water changes. At least 351 larvae were measured on day 12 from each treatment with five replicates each. Control larvae that did not receive any probiotic treatment ("Larvae only") had an average shell size of 153.21 µm, which was significantly smaller than that of the probiotic-supplemented larvae (Dunnett's Test, $P \leq .001$) (Table 1). Larvae from the treatment group "PB Multiple" that received repetitive probiotic doses with each water change had an average shell size of 161.24 µm, and the treatment group "PB Single" that received a single probiotic addition 24 hpf resulted in a statistically similar ($P = .26$), though a slightly larger average size of 165.49 µm (Table 1). Different letters denote statistical differences ($P \leq .001$). The boxes indicate the upper and lower quartiles, and the bar represents the median or middle quartile. The ends of the whiskers represent the most extreme values within the 1.5x interquartile range (IQR), and the empty circles indicate outliers.

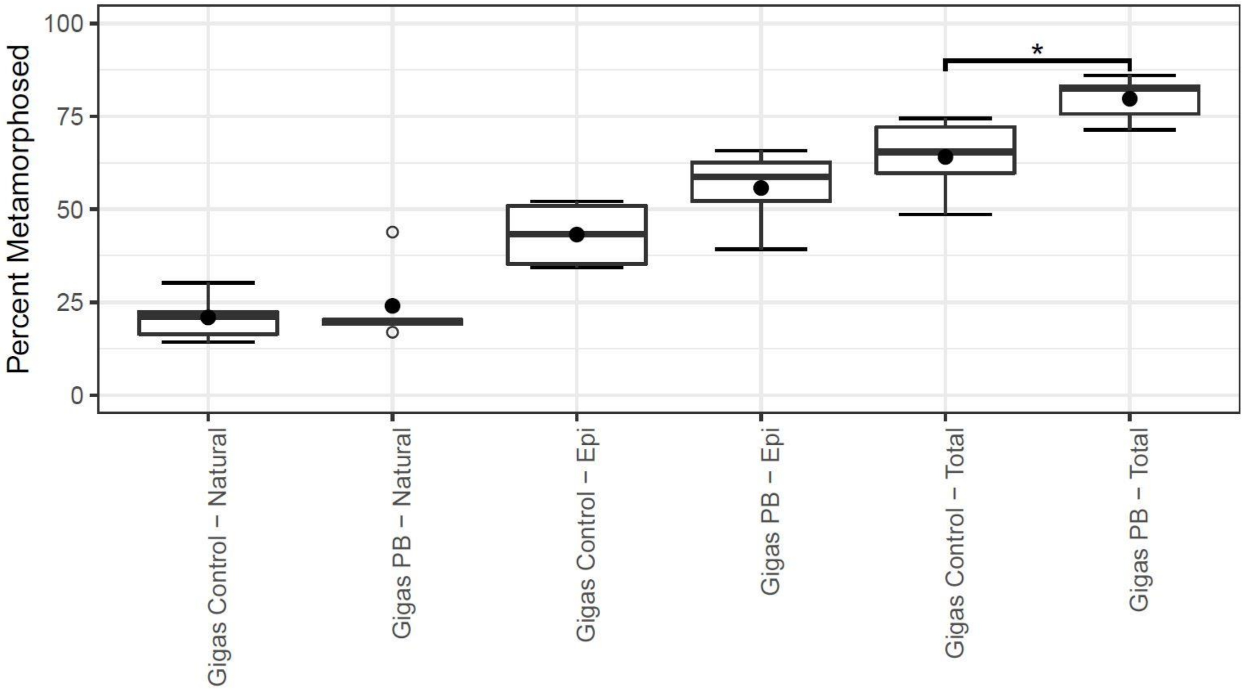


**S4 Figure: Improved total metamorphosis with a single probiotic addition for larvae of the Miyagi stock of *C. gigas* (*P*=.020*).** Larvae were first set naturally on marble tiles, then the remaining larvae were epinephrine (Epi) treated. “Total” indicates the mathematic sum of metamorphosed larvae that were naturally set or epinephrine-treated. Filled circles depict the average relative percent survival (RPS) of five replicate containers. The boxes indicate the upper and lower quartiles, and the bar represents the median or middle quartile. The ends of the whiskers represent the most extreme values within the 1.5x interquartile range (IQR), and the empty circles indicate outliers.

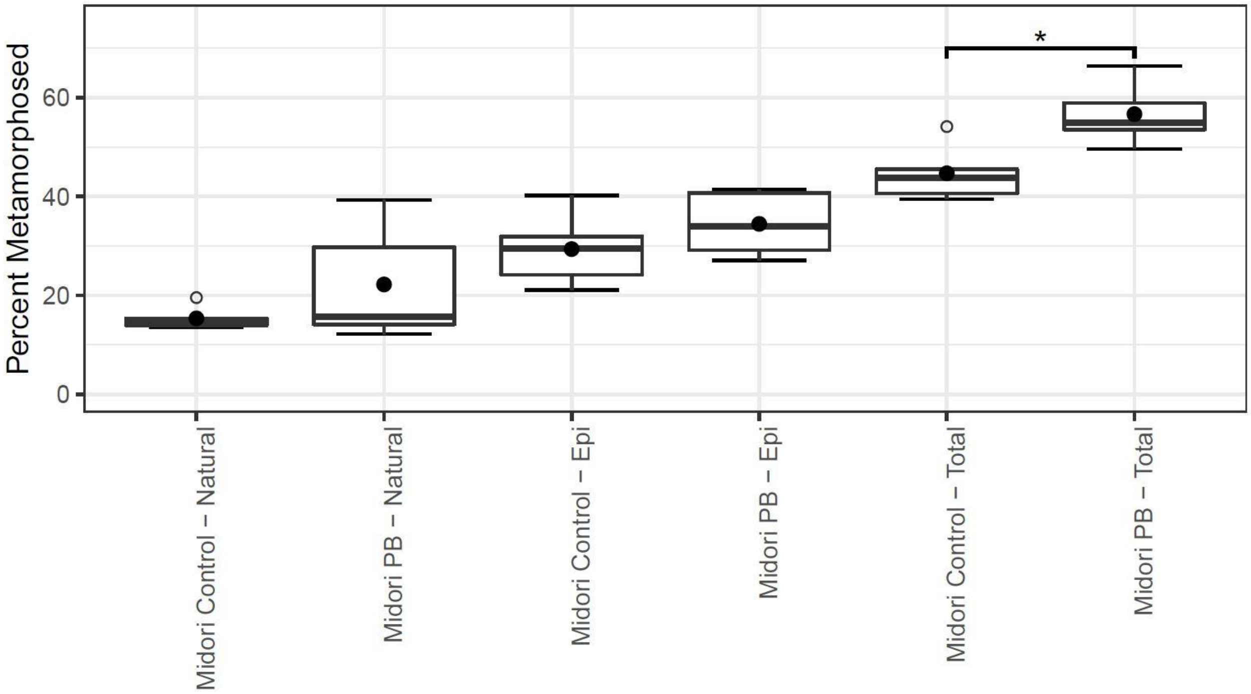


**S5 Figure: Improved total metamorphosis with a single probiotic addition for larvae of the Midori stock of *C. gigas* (*P*=.015*).** Larvae were first set naturally on marble tiles, then the remaining larvae were epinephrine (Epi)-treated. "Total" indicates the mathematic sum of metamorphosed larvae that were naturally set or epinephrine-treated. Filled circles depict the average relative percent survival (RPS) of five replicate containers. The boxes indicate the upper and lower quartiles, and the bar represents the median or middle quartile. The ends of the whiskers represent the most extreme values within the 1.5x interquartile range (IQR), and the empty circles indicate outliers.

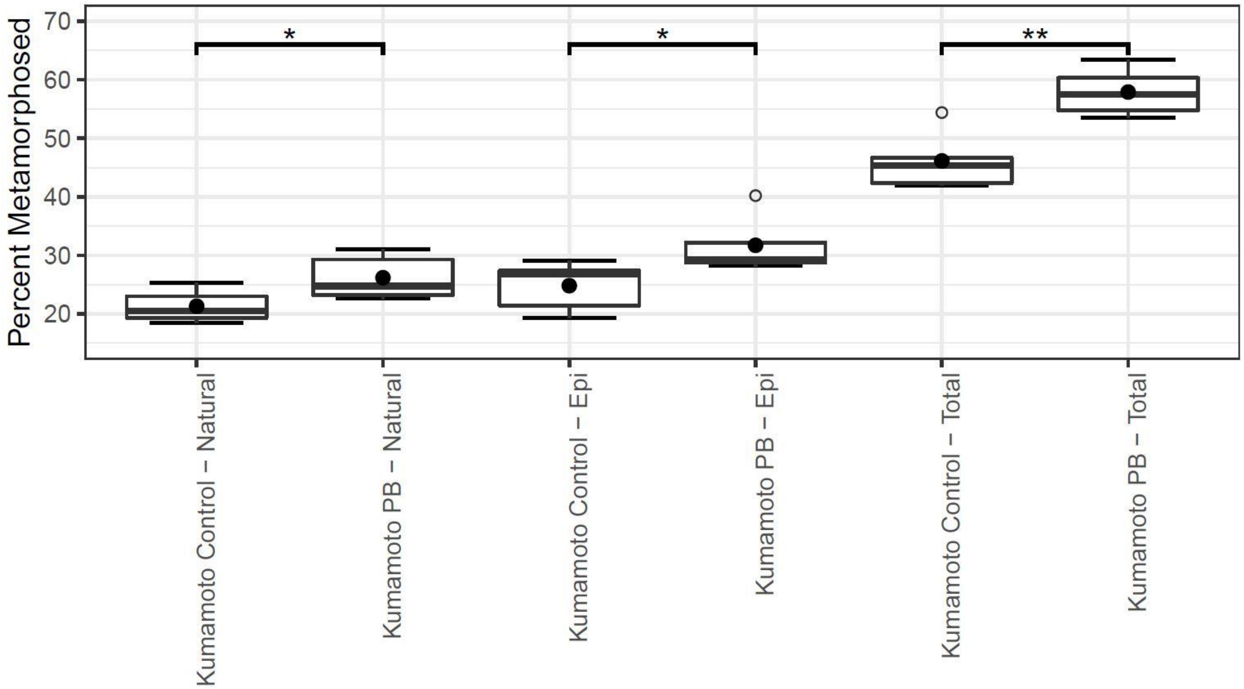


**S6 Figure: Improved natural (*P*=0.050*) and epinephrine-induced (*P*=.046*) metamorphosis with a single probiotic addition for larvae of Kumamoto oysters (*C. sikamea*) (*P*=.004**).** Larvae were first set naturally on marble tiles, then the remaining larvae were epinephrine (Epi) treated. “Total” indicates the mathematic sum of metamorphosed larvae that were naturally set or epinephrine-treated. Filled circles depict the average relative percent survival of five replicate containers. The boxes indicate the upper and lower quartiles, and the bar represents the median or middle quartile. The ends of the whiskers represent the most extreme values within the 1.5x interquartile range (IQR), and the empty circles indicate outliers.
