## Appendix Supplemental Tables for "A marine probiotic treatment against the bacterial pathogen *Vibrio coralliilyticus* to improve the performance of Pacific (*Crassostrea gigas*) and Kumamoto (*C. sikamea*) oyster larvae"

David Madison[a1], Carla Schubiger[b], Spencer Lunda[b], Ryan S. Mueller[c], Chris Langdon[a]

[a]Coastal Oregon Marine Experiment Station and Department of Fisheries, Wildlife, and Conservation Sciences, College of Agricultural Sciences, Oregon State University, Corvallis, Oregon, 97330, USA

[b]Department of Biomedical Sciences, Carlson College of Veterinary Medicine, Oregon State University, Corvallis, Oregon, 97330, USA

[c]Department of Microbiology, College of Science, Oregon State University, Corvallis, Oregon, 97330, USA

---

[1] Current address: JL. Trans Balauring, Desa Merdeka, Kecamatan Lebatukan, Kab. Lembata, 86681 NTT, Indonesia

### Appendix B. Supplemental tables.

**S1 Table: Pathogenicity testing of all 13 putative probiotic candidates with *C. gigas* larvae at 24 hpf.** Candidates (**bold**) that resulted in significantly reduced survival ($p \leq 0.05$) compared to that of the control (Larvae Only) were excluded from further testing.

| | | | | Dunn's Test (Benjamini-Hochberg Correction Applied) | | |
|---|---|---|---|---|---|---|
| **Treatment** | **Mean Survival** | **Std. Dev.[a]** | **SEM[b]** | **Comparison** | **Difference** | ***p*-Value** |
| B1 | 99.32 | 1.05 | 0.43 | B1 - Larvae Only | -0.41 | .4694 |
| B2 | 99.64 | 0.89 | 0.36 | B2 - Larvae Only | -0.09 | .492 |
| B3 | 100.00 | 0.00 | 0.00 | B3 - Larvae Only | 0.27 | .4694 |
| B8 | 99.35 | 1.60 | 0.65 | B8 - Larvae Only | -0.38 | .492 |
| B9 | 99.69 | 0.77 | 0.31 | B9 - Larvae Only | -0.05 | .492 |
| B10 | 100.00 | 0.00 | 0.00 | B10 - Larvae Only | 0.27 | .4694 |
| B11 | 99.40 | 0.95 | 0.39 | B11 - Larvae Only | -0.33 | .4694 |
| DM7 | 0.00 | 0.00 | 0.00 | DM7 - Larvae Only | -99.73 | **.0006*** ** |
| DM14 | 98.92 | 1.68 | 0.69 | DM14 - Larvae Only | -0.81 | .4694 |
| DM36 | 0.00 | 0.00 | 0.00 | DM36 - Larvae Only | -99.73 | **.000613*** ** |
| DM39 | 3.78 | 2.64 | 1.08 | DM39 - Larvae Only | -95.95 | **.00768** ** |
| D16 | 100.00 | 0.00 | 0.00 | D16 - Larvae Only | 0.27 | .4694 |
| D20 | 0.00 | 0.00 | 0.00 | D20 - Larvae Only | -99.73 | **.000613*** ** |
| Larvae Only | 99.73 | 0.66 | 0.27 | | | |

[a]Std. Dev. Indicates Standard Deviation; [b]SEM indicates Standard Error of the Mean; ** indicates $P \leq .01$; *** indicates $P \leq .001$

**S2 Table: Statistical comparison of the increased mean relative percent survival (RPS) of *C. gigas* larvae when treated with nine probiotic candidates compared to the positive control without probiotics (Vcor only) when exposed to *V. coralliilyticus* strain RE22.**

| | | | | **Dunn's Test (Benjamini-Hochberg Correction Applied)** | | |
|---|---|---|---|---|---|---|
| **Treatment** | **Mean RPS (%)** | **Std Dev[a]** | **SEM[b]** | **Comparison** | **Difference** | ***p*-value** |
| DM14 + Vcor | 99.71 | 0.87 | 0.36 | DM14 + Vcor – Vcor Only | **96.31** | **.0195*** |
| B1 + Vcor | 96.29 | 3.30 | 1.35 | B1 + Vcor – Vcor Only | **92.89** | **.04989*** |
| D16 + Vcor | 87.28 | 12.96 | 5.29 | D16 + Vcor – Vcor Only | **83.88** | .0766 |
| B11 + Vcor | 71.96 | 35.10 | 14.33 | B11 + Vcor – Vcor Only | **68.56** | .0818 |
| B2 + Vcor | 20.32 | 37.53 | 15.32 | B2 + Vcor – Vcor Only | 16.92 | .3798 |
| B8 + Vcor | 4.95 | 8.74 | 3.57 | B8 + Vcor – Vcor Only | 1.54 | .3545 |
| B3 + Vcor | 1.85 | 1.82 | 0.74 | B3 + Vcor – Vcor Only | -1.55 | .4898 |
| B9 + Vcor | 0.00 | 0.00 | 0.00 | B9 + Vcor – Vcor Only | -3.4 | .1097 |
| B10 + Vcor | 0.00 | 0.00 | 0.00 | B10 + Vcor – Vcor Only | -3.4 | .1097 |
| Vcor Only | 3.40 | 5.58 | 2.28 | | | |

[a]Std. Dev. indicates Standard Deviation; [b]SEM means Standard Error of the Mean; * indicates *P*≤.05

**S3 Table: Variability between replicate wells of the same treatments.** Variabilities of Relative Percent Survival (RPS) were most pronounced among wells of the treatments “Vcor Only”, “B11+Vcor”, and “D16+Vcor”.

| Treatment | RPS | Treatment | RPS | Treatment | RPS | Treatment | RPS | Treatment | RPS |
|---|---|---|---|---|---|---|---|---|---|
| Vcor Only | 0.00 | B1+Vcor | 92.08 | B11+Vcor | 8.53 | D16+Vcor | 91.36 | DM14+Vcor | 98.60 |
| Vcor Only | 14.6 | B1+Vcor | 94.13 | B11+Vcor | 56.56 | D16+Vcor | 100.3 | DM14+Vcor | 98.57 |
| Vcor Only | 1.97 | B1+Vcor | 98.91 | B11+Vcor | 75.68 | D16+Vcor | 70.19 | DM14+Vcor | 100.3 |
| Vcor Only | 1.89 | B1+Vcor | 100.3 | B11+Vcor | 97.04 | D16+Vcor | 92.56 | DM14+Vcor | 100.3 |
| Vcor Only | 0.00 | B1+Vcor | 94.00 | B11+Vcor | 95.57 | D16+Vcor | 97.32 | DM14+Vcor | 100.3 |
| Vcor Only | 1.93 | B1+Vcor | 98.34 | B11+Vcor | 98.41 | D16+Vcor | 71.99 | DM14+Vcor | 100.3 |

**S5 Table: Statistical comparison of different probiotic combination treatments shown in Figure 2.**

| Treatment | Mean RPS[a] | Std. Dev.[b] | SEM[c] | Diff.[d] | *p*-Value |
|---|---|---|---|---|---|
| D16 + Vcor | 15.24 | 17.02 | 6.95 | 9.01 | .996 |
| B11 + Vcor | 28.98 | 32.83 | 13.40 | 22.74 | .488 |
| DM14 + Vcor | 65.17 | 28.91 | 11.80 | 58.93 | **.003**** |
| B11 D16 + Vcor | 70.40 | 35.17 | 14.36 | 64.17 | **.001***** |
| DM14 D16 + Vcor | 52.28 | 21.35 | 8.72 | 46.05 | **.034*** |
| B11 DM14 + Vcor | 85.88 | 26.82 | 10.95 | 79.65 | **<.001***** |
| B11 DM14 D16 + Vcor | 71.18 | 30.72 | 12.54 | 64.95 | **.001***** |
| Vcor Only | 6.23 | 4.01 | 1.64 | | |

[a]RPS indicates Relative Percent Survival; [b]Std. Dev. indicates Standard Deviation; [c]SEM indicates Standard Error of the Mean; [d]Diff. indicates Difference to Vcor Only control; * indicates $P \leq .05$; ** indicates $P \leq .01$; *** indicates $P \leq .001$

**S6 Table: Statistical comparison of different probiotic combination treatments.**

| Treatment | Mean RPS[a] | Standard Deviation | SEM[b] |
|---|---|---|---|
| 1 Strain | 36.46 | 33.43 | 7.88 |
| 2 Strains | 69.52 | 30.15 | 7.11 |
| 3 Strains | 71.18 | 30.72 | 12.54 |
| Vcor Only | 6.23 | 4.01 | 1.64 |

| Dunn's Test (Benjamini-Hochberg Correction Applied) - Pairwise comparisons | Difference | *p*-value |
|---|---|---|
| 1 Strain - Vcor Only | 30.23 | **.0377*** |
| 2 Strains - Vcor Only | 63.29 | **.0003***** |
| 3 Strains - Vcor Only | 64.95 | **.0024**** |
| 1 Strain - 2 Strains | -33.06 | **.0071**** |
| 1 Strain - 3 Strains | -34.72 | **.0339*** |
| 2 Strains - 3 Strains | -1.66 | .4983 |

[a]RPS indicates Relative Percent Survival; [b]SEM indicates Standard Error of the Mean; * indicates $P<.05$; ** indicates $P<.01$; *** indicates $P<.001$

**S7 Table: Single probiotic application increased successful metamorphosis of *C. gigas* (Miyagi stock) larvae.**

| Treatment | Mean % Natural Set | Mean % Epi Set | Mean % Total Set | SD Natural Set | SEM Natural Set | SD Epi Set | SEM Epi Set | SD Total Set | SEM Total Set |
|---|---|---|---|---|---|---|---|---|---|
| Larvae Only | 2.24 | 4.81 | 7.06 | 1.72 | 0.77 | 7.02 | 3.14 | 8.66 | 3.87 |
| Probiotics Once | 8.29 | 13.03 | 21.32 | 2.07 | 0.93 | 3.14 | 1.41 | 4.65 | 2.08 |
| Probiotics Repeated | 2.80 | 4.24 | 7.04 | 0.77 | 0.35 | 2.13 | 0.95 | 2.67 | 1.19 |

| | **Dunnett's Test (Comparing to Larvae Only Control)** | | | |
|---|---|---|---|---|
| | **Difference** | **95% CI** | **99% CI** | ***p*-Value** |
| Probiotics Once | 14.27 | (4.96, 23.58) | (1.66, 26.87) | **.004**** |
| Probiotics Repeated | -0.01 | (-9.32, 9.30) | (-12.62, 12.59) | >.999 |

** indicates $P \leq .01$

**S8 Table: Two-sample t-tests comparing natural, epinephrine-induced, and total metamorphosis and settlement with a single probiotic treatment versus a non-probiotic control for of *C. gigas* (Miyagi and Midori stocks) and *C. sikamea* (Kumamoto) larvae.**

| **NATURAL** | Comparison | **Two Sample t-test - Natural** | | | |
|---|---|---|---|---|---|
| | | **Difference** | **95% CI** | **99% CI** | ***p*-Value** |
| Miyagi | Control - PB | -3.12 | (-16.29, 10.05) | (-22.28, 16.04) | .599 |
| Midori | Control - PB | -6.87 | (-19.32, 5.57) | (-24.98, 11.24) | .239 |
| Kumamoto | Control - PB | -4.87 | (-9.74, 0.01) | (-11.96, 2.23) | **.0503*** |
| **EPINEPHRINE** | Comparison | **Two Sample t-test - Epi** | | | |
| | | **Difference** | **95% CI** | **99% CI** | ***p*-Value** |
| Miyagi | Control - PB | -12.53 | (-26.39, 1.33) | (-32.69, 7.64) | .071 |
| Midori | Control - PB | -5.10 | (-15.30, 5.10) | (-19.94, 9.74) | .282 |
| Kumamoto | Control - PB | -6.90 | (-13.63, -0.17) | (-16.69, 2.89) | **.04564*** |
| **TOTAL** | Comparison | **Two Sample t-test - Total** | | | |
| | | **Difference** | **95% CI** | **99% CI** | ***p*-Value** |
| Miyagi | Control - PB | -15.65 | (-28.07, -3.24) | (-33.71, 2.41) | **.01965*** |
| Midori | Control - PB | -11.97 | (-20.86, -3.08) | (-24.90, 0.96) | **.01454*** |
| Kumamoto | Control - PB | -11.76 | (-18.42, -5.11) | (-21.45, -2.08) | **.00355**** |

* indicates $P \leq .05$; ** indicates $P \leq .01$
